# A myosin hypertrophic cardiomyopathy mutation disrupts the super-relaxed state and boosts contractility by enhanced actin attachment

**DOI:** 10.1101/2025.06.02.657466

**Authors:** Robert C. Cail, Bipasha Barua, Faviolla A. Báez-Cruz, Donald A. Winkelmann, Yale E. Goldman, E. Michael Ostap

## Abstract

Hypertrophic cardiomyopathy (HCM) is a leading cause of cardiac failure among individuals under 35. Many genetic mutations that cause HCM enhance ventricular systolic function, suggesting that these HCM mutations are hypercontractile. Among the most common causes of HCM are mutations in the gene MYH7, which encodes for β-cardiac myosin, the principal human ventricular myosin. Previous work has demonstrated that, for purified myosins, some MYH7 mutations are gain-of-function while others cause reduced function, so how they lead to enhanced contractility is not clear. Here, we have characterized the mechanics and kinetics of the severe HCM-causing mutation M493I. Motility assays demonstrate a 70% reduction of actin filament gliding velocities on M493I-coated surfaces relative to WT. This mutation slows ADP release from actomyosin·ADP 5-fold without affecting phosphate release or ATP binding. Yet it enhances steady-state ATPase *V_max_* 2-fold. Through single-molecule mechanical studies, we find that M493I myosin has a normal working stroke of 5 nm but a significantly prolonged actin attachment duration. Under isometric feedback, M493I myosins produce high, sustained force, with an actin detachment rate that is less sensitive to force than that of WT myosin. We also report direct measurement of the equilibrium state of the super-relaxed to disordered relaxed (SRX-DRX) regulatory transition and show its disruption in M493I, with a concomitant enhancement to actin attachment kinetics. Together, these data demonstrate that enhanced myosin binding from inhibition of myosin’s off state, combined with slow ADP release and enhanced force production, underlie the enhanced function and etiology of this HCM mutation.

**Significance Statement:** Hypertrophic cardiomyopathy (HCM) is a leading genetic cause of sudden cardiac death in young individuals. Although often described as a hypercontractile disease, the molecular basis for this remains unclear, especially for mutations with inhibitory effects in various *in vitro* assays. We show that the severe HCM mutation M493I in β-cardiac myosin slows ADP release yet enhances force output and actin attachment through multiple mechanisms, including disrupted autoinhibition via the super-relaxed state. Our findings unify seemingly contradictory biophysical changes into a coherent mechanistic model and support the hypothesis that increased myosin head availability, rather than enhanced individual kinetics alone, underlies HCM hypercontractility.

## Introduction

Hypertrophic cardiomyopathy (HCM) is an inherited genetic disorder that follows an autosomal dominant pattern and affects approximately 1 in 500 individuals (1). It is a leading cause of heart failure and sudden cardiac death, particularly in individuals under 35 (2). HCM is characterized by an abnormal thickening of the left ventricular free wall and intraventricular septum, along with cardiomyocyte disarray and interstitial fibrosis (3–6). These changes can lead to complications such as outflow obstruction, atrial fibrillation, heart failure, ventricular arrhythmias, and sudden cardiac death (1–8).

HCM-causing mutations can occur in approximately 20 sarcomeric genes, with mutations in the gene MYH7, encoding β-cardiac myosin, the principal paralog of myosin found in human ventricles, being implicated in 30–40% of cases (9, 10). The majority of these are classified as missense mutations (11). HCM is typically associated with impaired diastolic function while systolic function is either preserved or enhanced, leading to its classification as a hypercontractile disorder (12–14). However, mutations in β-cardiac myosin (hereafter, myosin) can either increase or decrease force production when analyzed in isolated myofibrils or single-molecule studies (15–19). It has been unclear how these opposing effects result in the same disease phenotype.

One HCM-causing mutation, M493I, is linked to septal stiffening, severe physical limitations, and congestive heart failure and sudden cardiac death (20, 21). The M493 residue is located in the relay helix between the active site and the rotating converter domain of β-cardiac myosin, where it may form a hydrogen bond with C705 in the SH1 helix (22–26) (Fig 1A). The relay helix plays a crucial role in transmitting the nucleotide state of the active site to the converter domain, coupling ATP hydrolysis and product release to the tilting of myosin’s lever arm (27, 28). It is highly conserved across myosin paralogs (Fig S1A) (29). While M493 is conserved across fast class II myosins in humans, other myosin paralogs have different amino acids in this position, such as isoleucine (class VI and IX myosins) or valine (class V and VII myosins). The region near M493 is a hotspot for HCM- and dilated cardiomyopathy (DCM)-causing mutations, including residues E497, Y501, and F513 in the relay helix, as well as R712 and F764 in the converter domain (16, 23–26).

**Fig 1:**
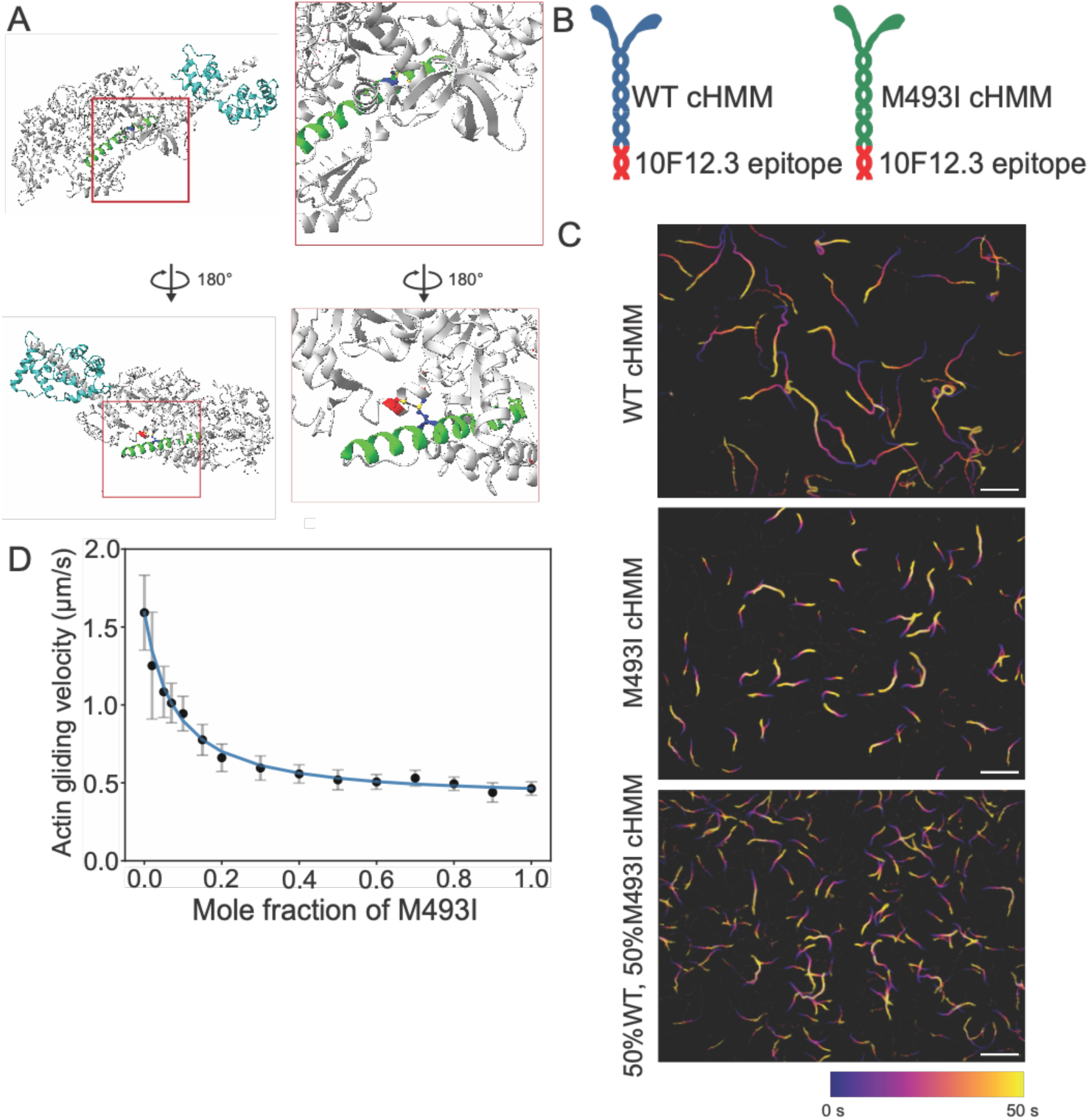
M493I is a relay helix mutation that significantly slows actin gliding velocity. A) The relay helix (green) contains M493 (blue), which establishes a hydrogen bond with C705 (red) in the converter domain. B) Schematic of purified WT- and M493I-cHMMs, containing 42 heptads of coiled-coil tail and engineered with a coiled-coil epitope (10F12.3 subfragment of G. domesticus skeletal muscle myosin). C) Color-coded time projections from actin gliding assays WT (top), M493I (middle) and 50-50 mixture of the two myosins (bottom). Longer traces correspond to faster gliding velocity. Scale bars: 10 μm. D) Actin gliding velocity as a function of mole fraction of M493I, demonstrating decreased gliding speeds with low proportions of M493I motors present.

In addition to the canonical actin-activated ATPase mechanism (30), many class II myosins, including β-cardiac myosin, have a conserved “off” state that prevents actin binding (31–33). This state is stabilized by an interaction between the motor domain (head) and the dimer- and filament-forming tail domain, and it is regulated by phosphorylation and other factors. In muscle myosins, this state is referred to as the interacting-head motif (IHM) and is linked to the super-relaxed (SRX) state of myosin (34–36). The SRX state is thought to consume ATP at a 10-fold slower rate than the alternative disordered relaxed (DRX) state, in which the myosin heads are freer and more available for actin binding. The mechanisms and kinetics governing the SRX/DRX transition for purified heavy meromyosin (HMM) remain unclear. However, structural studies have shown that the IHM state corresponds to the pre-power-stroke (PPS) conformation, with ADP·P_i_ bound in the active site (37).

Efforts to develop a unifying model of HCM have led to the “head availability hypothesis” (14, 38–41). This hypothesis suggests that HCM-causing mutations disrupt the SRX/DRX balance, increasing the number of myosin heads in the DRX (available) state. This increase in available heads causes excessive activation of the thin filament and hypercontractility, regardless of the mutation’s direct effects on intrinsic myosin kinetics, working stroke size, and power output.

In this study, we characterized the impact of the M493I mutation on the mechanochemistry of a recombinant HMM construct. We applied ensemble kinetics and the single-molecule optical trapping technique to measure the effect of the mutation on key ATPase cycle transitions, linking these changes to specific substeps of energy transduction by myosin. Our findings provide evidence that the SRX/DRX equilibrium is fast relative to ATP turnover, along with an estimation of its equilibrium constant, insights into how the M493I mutation alters head availability in myocytes, and a measurement of the enhanced actin-binding activity at the single-molecule level induced by M493I. Together, these results point to a mechanism by which a single HCM-linked mutation alters myosin’s mechanochemical cycle to promote hypercontractility.

## Results

### M493I slows actin gliding velocity

Human β-cardiac myosin wild type heavy meromyosin (WT-cHMM) and M493I mutant (M493I-cHMM) constructs, with 42 heptads of the coiled-coil S2 tail domain, were expressed in mouse C2C12 myoblasts and purified (Fig S1B) (16, 25). At the C-terminus of the S2 domain, a coiled-coil epitope tag comprised of the 10F12.3 region of *Gallus domesticus* skeletal muscle myosin was included, allowing for site-specific binding of purified myosins through a monoclonal antibody to this epitope tag (Fig 1B) (42).

To assess whether the M493I mutation affects unloaded actin motility, we conducted an *in vitro* actin gliding assay using both WT and M493I myosins. Adhering the myosin constructs via an antibody to the 10F12.3 motif enabled precise deposition of mixed mole fractions of WT and M493I myosins onto the coverslips at a total loading concentration of 10 µg/mL. WT myosin exhibited persistent actin propulsion, with a velocity of 1.6 ± 0.2 µm/s (Fig. 1C-D, Supplemental Video 1, Table 1). M493I myosin also supported smooth continuous actin gliding but at a 72% slower velocity (0.46 ± 0.04 µm/s; Fig. 1C-D, Supplemental Video 2, Table 1). When WT and M493I myosins were co-incubated in varying proportions, actin gliding velocity declined sharply with increasing M493I mole fraction, showing a 50% velocity reduction at just 10% proportion of mutant myosin and approaching the low-velocity asymptote of the velocity curve at ∼50% M493I myosin (Fig. 1C-D, Supplemental Video 3, Table 1). These data were fitted to a quadratic model of loaded force production in the presence of mixed myosin species, as described previously (see Methods) (43, 44). The concave upward shape of this curve suggests that the faster WT-cHMM experiences frictional loading by the slower M493I-cHMM, indicative of drag due to enhanced actin attachment of the mutant version (44). To further investigate this frictional slowing, we examined which steps of the mechanochemical cycle were affected by the M493I mutation.

**Table 1:** Kinetic and mechanical parameters of WT-cHMM and M493I-cHMM.

|  | WT-cHMM | M493I-cHMM | Method |
| --- | --- | --- | --- |
| **Actin gliding velocity ($\mu\text{m/s}$)** | $1.6 \pm 0.2$ | $0.46 \pm 0.04$ | In vitro actin gliding velocity with rhodamine-phalloidin labelled actin |
| **P<sub>i</sub> release rate (fast phase) ($\text{s}^{-1}$)** | $15.8 \pm 5.0$ | $15.1 \pm 4.0$ | Phosphate-binding protein fluorescence, SF, 30 $\mu\text{M}$ porcine TFs |
| **P<sub>i</sub> release fast amplitude** | 0.27 | 1 | Phosphate-binding protein fluorescence, SF, 30 $\mu\text{M}$ porcine TFs |
| **P<sub>i</sub> release rate (slow phase) ($\text{s}^{-1}$)** | $0.37 \pm 0.08$ | N/A | Phosphate-binding protein fluorescence, SF, 30 $\mu\text{M}$ porcine TFs |
| **ADP release rate ($\text{s}^{-1}$)** | $69 \pm 7$ | $12.0 \pm 0.8$ | Pyrene-actin, SF |
| **ATP binding second order rate ($\mu\text{M}^{-1}\text{s}^{-1}$)** | 4.7<br>4.6-5.2 95% CI | 6.2<br>5.8-6.4 95% CI | Pyrene-actin, SF |
| **Maximum rate ATP-induced dissociation of actomyosin** | $1191 \pm 109^*$<br>$941 \pm 60^{**}$ | 340<br>322-419 95% CI | Pyrene-actin, SF |
| **ATP binding $K_M$ ($\mu\text{M}$)** | $128 \pm 29^{**}$ | 35.4<br>24.7-61.8 95% CI | Pyrene-actin, SF |
| **Steady-state ATPase $V_{\text{max}}$ ($\text{s}^{-1}$)** | 1.20<br>1.09-1.77 95% CI | 2.35<br>1.51-2.51 95% CI | NADH-coupled ATPase assay, spectrophotometer, porcine TFs |
| **Steady-state ATPase $K_M$ ($\mu\text{M}$)** | 6.66<br>5.25-13.62 95% CI | 7.65<br>2.97-10.09 95% CI | NADH-coupled ATPase assay, spectrophotometer, porcine TFs |
| **$k_{\text{detach}}$ (1 $\mu\text{M}$ ATP) ($\text{s}^{-1}$)** | 6.09<br>5.66-6.59 95% CI | 6.26<br>5.89-6.68 95%CI | 3-bead optical trap assay |
| **$k_{\text{detach}}$ (2 mM ATP) ($\text{s}^{-1}$)** | 58.48<br>51.95-67.02 95%CI | 12.06<br>11.11-13.26 95%CI | 3-bead optical trap assay |
| **Substep 1 (nm)** | $3.88 \pm 0.23$ (SEM) | $3.80 \pm 0.22$ (SEM) | 3-bead optical trap assay |
| **Substep 2 (nm)** | $0.94 \pm 0.18$ (SEM) | $0.88 \pm 0.15$ (SEM) | 3-bead optical trap assay |
| **Total step size (*nm*)** | 4.82 ± 0.25 (SEM) | 4.68 ± 0.21 (SEM) | 3-bead optical trap assay |
| **Bell equation $k_0$ ($s^{-1}$)** | 32.9<br>28.8-35.3 95% CI | 12.9<br>9.2-18.2 95%CI | 3-bead optical trap assay, isometric FB |
| **Bell equation distance parameter (*nm*)** | 0.80<br>0.62-0.97 95% CI | 0.32<br>0.00-0.70 95% CI | 3-bead optical trap assay, isometric FB |
| **Single-nucleotide turnover rate (NT) ($s^{-1}$)** | 0.0047 ± 0.0005*** | 0.0086 ± 0.0001 | mantATP turnover, SF |
| **Single-nucleotide turnover rate (undigested) ($s^{-1}$)** | 0.0042 ± 0.0004*** | 0.0065 ± 0.0003 | mantATP turnover, SF |
| **Single-nucleotide turnover rate (papain) ($s^{-1}$)** | 0.013 ± 0.001*** | 0.014 ± 0.006 | mantATP turnover, SF |
| **SRX-DRX equilibrium constant** | 0.32 ± 0.03*** | 0.63 ± 0.08 | mantATP turnover, SF |
| **Actin re-attachment rate in optical trap (slow) ($s^{-1}$)** | 0.83<br>0.70-0.9 95% CI | 1.62<br>1.45-1.74 95% CI | 3-bead optical trap assay, pedestal bead stage feedback |
| **Actin re-attachment slow amplitude** | 80%<br>65-87 95% CI | 81%<br>71-85 95% CI | 3-bead optical trap assay, pedestal bead stage feedback |
| **Actin re-attachment rate in optical trap (fast) ($s^{-1}$)** | 17<br>5.5-36 95% CI | 23<br>11-39 95% CI | 3-bead optical trap assay, pedestal bead stage feedback |

Table Legend: Experiments performed in KMg25 buffer.
\*Published previously (16)
\*\*Published previously (46)
\*\*\*Published previously (60)

### M493I preserves P_i_ release and ATP binding but slows ADP release

We investigated whether the M493I mutation alters key rate constants in the ATPase cycle that regulate entry into and exit from the force-bearing, actin-bound states using stopped-flow kinetics (45). The rate of the phosphate release step, which limits entry into the strong-binding states of the ATPase cycle, was measured by fluorescent Pi binding protein (46) in the presence of a saturating concentrations (30 µM) of porcine ventricular thin filaments (Fig. S2A). As found previously (47, 48), P_i_ release transients in the presence of WT-cHMM were best fit by a two-exponential function, with a fast-phase (15.8 ± 5 s⁻¹, 27% of the transient amplitude, Fig. S2B, Table 1) and slow-phase (0.37 ± 0.08 s⁻¹, 73% of the transient amplitude, Fig. S2B, Table 1). M493I-cHMM released P_i_ in a single exponential phase at a rate of 15.1 ± 4 s⁻¹, which was similar to the fast-phase rate of WT-cHMM (Fig S2C, Table 1).

To evaluate exit from the force-bearing states, we measured the rate of ADP release and ATP binding to actin-attached myosin bound by dissociation from pyrene-actin (see Methods; Fig. S2D-E). ADP release from M493I-cHMM was approximately five-fold slower (12.0 ± 0.8 s⁻¹; Table 1) than from WT-cHMM (69 ± 7 s⁻¹; Table 1). ATP-induced dissociation of pyrene-actin-bound myosin was measured for a range of MgATP concentrations (500 nM – 0.6 mM). Pyrene fluorescence transients were well fit by single exponential functions (Fig S2 F-G). The apparent second-order rate constant for ATP binding to M493I-cHMM (6.2 (µM·s)^−1^; 5.8–6.4 (µM·s)^−1^ 95% confidence interval (CI)) was slightly faster than WT-cHMM (4.7 (µM·s)^−1^; 4.6–5.2 (µM·s)^−1^ 95% CI; Fig. S2H, Table 1). However, the maximum rate of ATP-induced dissociation for M493I-cHMM (340 s⁻¹; 322–419 s⁻¹ 95% CI) was slower than the previously determined rate for WT-cHMM (1191 ± 109 s⁻¹) (Fig S2I) (16). ADP release rather than ATP binding limits exit from the strong-binding states under mM physiological ATP concentrations. Thus, our findings suggest that the observed slowing of actin gliding is primarily attributable to the reduced rate of ADP release.

### M493I preserves the power stroke and enhances actin attachment duration

Some HCM-causing mutations in myosin can significantly alter the power stroke and actin filament sliding, particularly those near the lever arm (16, 17, 19). We examined the impact of the M493I mutation on myosin’s power stroke using optical trapping assays to assess how this relay-helix mutation affects motor properties and muscle physiology.

Pedestal beads, sparsely coated with antibodies against the 10F12.3 epitope tag, were used to bind WT-cHMM or M493I-cHMM, at sufficiently sparse coverage (with only one in five beads interacting with actin) to achieve single molecule events. An actin filament was suspended between two optically trapped beads, and bead position and variance were monitored. Single actomyosin binding events were detected by a decrease in covariance between the beads (Fig. 2A, 49, 50). For both WT and M493I myosins at 1 µM MgATP, transient decreases in covariance corresponded with bead displacement during attachment events (Fig. 2A). Interaction duration decreased with increasing ATP, following single-exponential cumulative distribution. The detachment rate (*k*_detach_) at 1 µM ATP was 6.09 s⁻¹ (5.66 - 6.59 s⁻¹ 95% CI) for WT and 6.26 s⁻¹ (5.89 - 6.68 s⁻¹ 95% CI) for M493I, consistent with detachment rate being limited by ATP binding (Fig. 2B-C). At 2 mM MgATP, actin detachment rate is limited by ADP release and M493I detached more slowly (*k*_detach_ = 12.06 s⁻¹, 11.11 - 13.26 s⁻¹ 95% CI) compared to WT (*k*_detach_ = 58.48 s⁻¹, 51.95 - 67.02 s⁻¹ 95% CI). This result demonstrates that the M493I mutation increases the duration of myosin attachments to actin, consistent with slowed ADP release which, in turn, explains its slowing of *in vitro* filament gliding (Fig. 2B-C).

**Fig 2:**
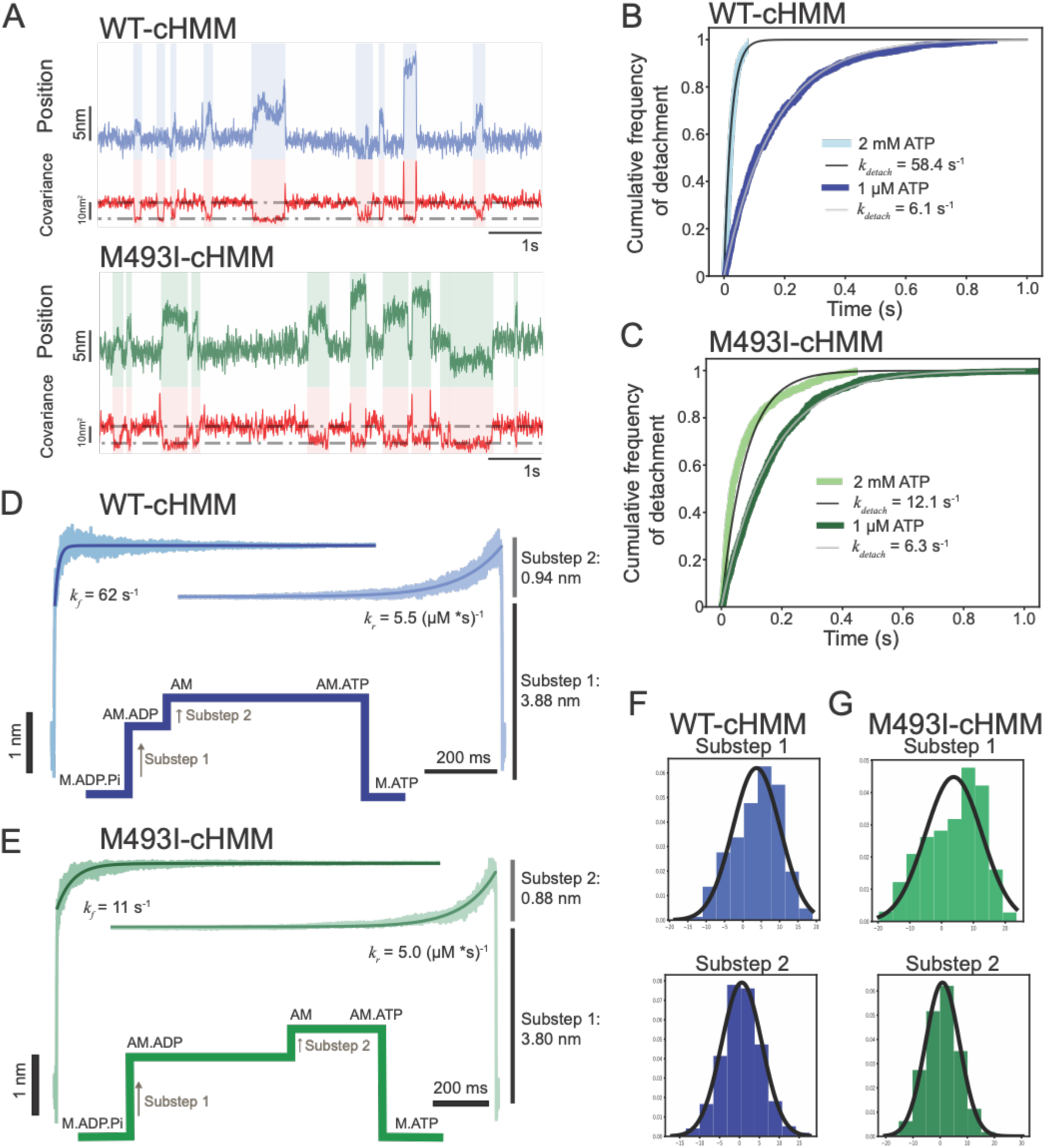
Step size and duration for WT and M493I myosins. A) Sample traces from unloaded optical trap assays of WT and M493I-cHMM. B-C) Cumulative distributions of attachment durations with fitted single-exponential curves showing the difference in attachment lifetime at high [ATP] between WT (B) and M493I-myosin (C). D-E) Ensemble averages and single-exponential fits of WT (D) and M493I (E) myosin events at 1 μM [ATP] demonstrating the same step amplitudes but slowed transition from substep 1 to 2 for M493I myosin. F-G) Average substep sizes for WT (F) and M493I (G) myosins with fitted Gaussian curves.

The cardiac myosin power stroke exhibits two barbed-end directed steps, linked to (1) P_i_ release from AM·ADP·P_i_, and (2) ADP release from AM·ADP. The average total amplitude of the working stroke was determined by combining single-molecule interactions aligned at initial attachment times (time-forward ensemble averages) and detachment times (time-reversed averages) (Fig. 2D-E) (51, 52). For time-forward averages, detected events were aligned at the start of the interaction, extending shorter events for averaging in the software to same duration as longest detected interaction for averaging. The exponential increase in these averages after the initial displacement is caused by the transition from substep 1 to substep 2, i.e. ADP release (52–54). For reverse averages, the ends of each interaction were aligned, extending each shorter interaction back in time to match the longest duration for averaging. The exponential increase in displacement in these averages reports the rate of ATP binding before detachment.

For wild-type myosin substeps 1 and 2 had displacements of 3.88 ± 0.23 nm (SEM) and 0.94 ± 0.18 nm (SEM) for a total step size of 4.82 ± 0.25 nm (SEM) (Fig 2D, Fig 2F). M493I-myosin’s ensemble averages a 1 µM ATP revealed a quite similar two-substep working stroke as that of WT-myosin with substep 1 and 2 displacements of 3.80 ± 0.22 nm (SEM) and 0.88 ± 0.15 nm (SEM) for a similar total step size of 4.68 ± 0.21 nm (SEM) (Fig 2D, Fig 2G).

For WT-myosin at 1 µM ATP, the ADP release rate and ATP binding rate determined by fitting the exponential phases of the forward and reverse ensemble averages, respectively, were 62 s^−1^ (58 – 71, 95% CI) and 5.5 s^−1^ (3.6 - 8.6, 95% CI) in good agreement with values from pyrene-actin bulk transients (Fig 2D). For M493I-myosin, the rate for transition from substep 1 to substep 2 was 11 s^−1^ (9.5-13.3, 95% CI) quite similar to the 12.0 s^−1^ ADP off-rate as measured from the bulk pyrene actin kinetics, but slower than WT. The rate constant of transition from substep 2 to unbound myosin, due to ATP binding was 5.0 s^−1^ (4.2-6.5, 95% CI) quite similar to WT and to the 6.2 (µM·s)^−1^ measured by pyrene actin (Fig 2E). Thus, M493I alters the working stroke of actomyosin not by altering the step size or ATP binding rate, but by slowing detachment through slowing the transition from substep 1 to substep 2, consistent with slowed ADP release and enhanced frictional loading of a moving actin filament.

### Under isometric tension, M493I produces remarkably long-duration, high-force actin attachments

Myosin’s loaded force generation capability, with associated load-dependent release of ADP release, is a key factor in effective, synchronized cardiac muscle shortening and energetic regulation (55–57). HCM-causing mutations in MYH7 can either increase or decrease the force produced by actomyosin interactions (16, 17, 19). We sought to investigate the effect of the mutation M493I on force production and load-dependent ADP dissociation kinetics using the three-bead optical trap assay under isometric feedback conditions (57). In this assay, the position of the bead at the pointed end of the actin dumbbell (termed the transducer bead) is monitored optically, and the position of the barbed-end bead (the motor bead) is adjusted by feedback to an electro-optic deflector to minimize (clamp) the displacement of the transducer bead, thus approximating the isometric condition in a muscle independent of series compliances in the dumbell. When myosin interacts with the actin filament, it is subjected to hindering load applied by this feedback as it produces its working stroke, and we can assay both its force generation capability and its loaded ADP dissociation rate.

WT-cHMM demonstrates regular interactions of variable force under load, as expected from the thermal (Brownian) fluctuations at the moment of attachment (Fig 3A, top) (54, 57). M493I-cHMM interactions under load are striking in their duration and force magnitude to a level that often exceeds the 25 pN dynamic range of our isometric clamp feedback loop (Fig 3A, bottom). Histograms of the forces from individual interactions demonstrate a considerable increase in the proportion of high-force interactions for M493I myosins relative to WT; the force measurements plotted are an underestimate because the highest force value is limited by the aforementioned dynamic range of the instrument (Fig 3B). We propose that these high forces are the result of recruitment of the second head of the HMM.

**Fig 3:**
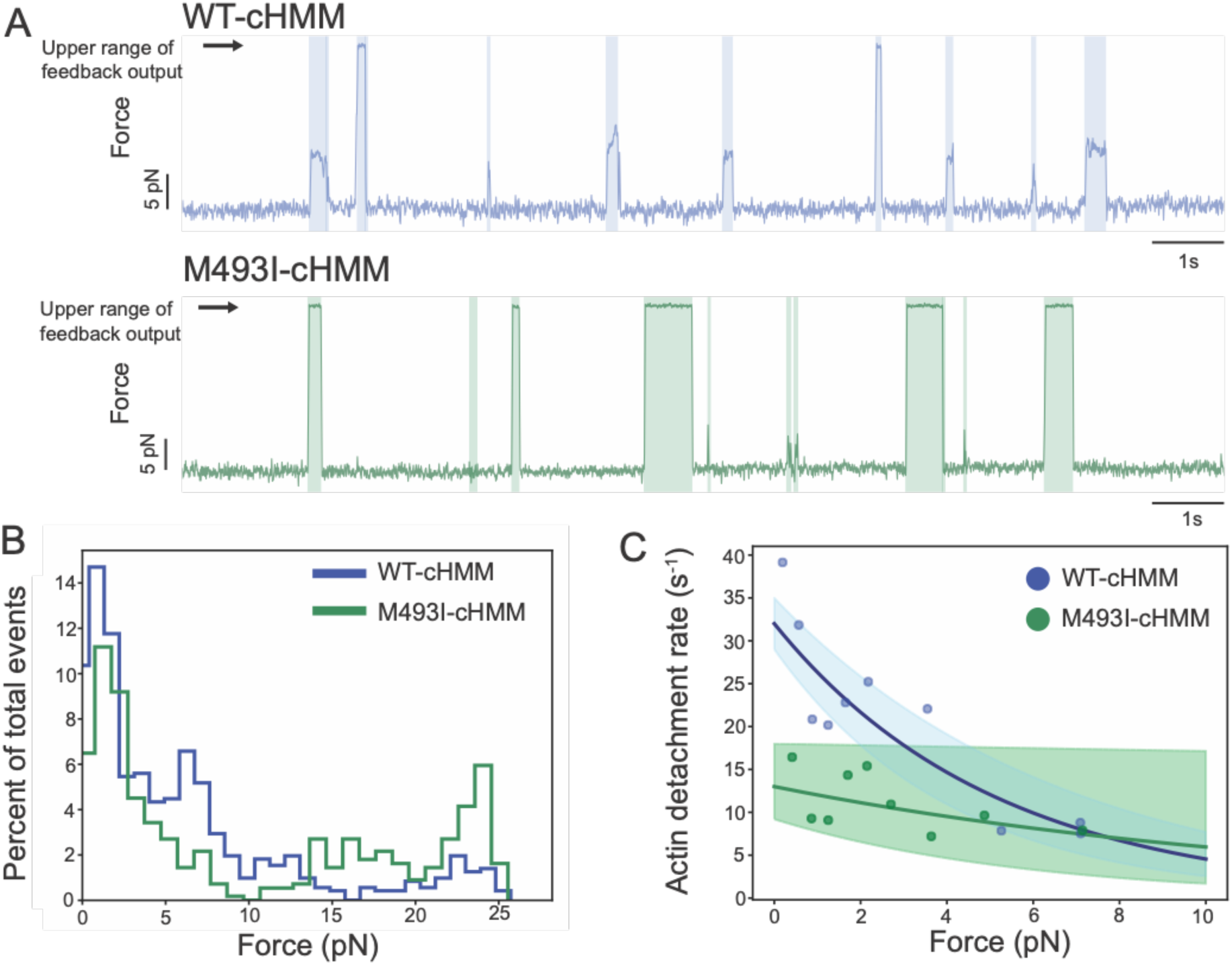
Force production by WT and M493I myosins. A) Sample traces from isometric feedback optical trap experiments showing a high proportion of feedback-saturating events present with M493I myosin. B) Histograms of force distribution for WT and M493I myosins. The mutant exhibits more high-force (>10 pN) events than WT. C) Actin detachment rate vs force for WT (blue) and M493I (green) myosins showing similar force-induced slowing of ADP release. Data are averaged in 25-point bins. Solid lines represent fits to the Bell equation applied to unbinned data, and shaded areas indicate the 95% confidence intervals, derived from bootstrap-based Bell equation fits (see Methods).

When interactions >10 pN are excluded from both datasets (to limit the distributions to interactions within the linear force range), the detachment rate *vs*. force data can be fitted to the Bell equation (58),

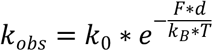

Where *k_obs_* is the observed detachment rate under load, *k*_0_ is the detachment rate (limited by ADP release) at zero force, *F* is the force applied to the myosin, *k_B_*T* is the Boltzmann constant times temperature, and *d* is the distance parameter, the effective displacement between the force-generating attached position and the transition barrier to detachment. WT *k_0_* = 32.0 s^−1^ (28.8 - 35.3 s^−1^ 95% CI) and *d* = 0.80 nm (0.62 - 0.97 nm 95% CI), whereas for M493I, *k_0_* = 12.9 s^−1^ (9.2 – 18.2 s^−1^ 95% CI) and *d* = 0.32 nm (0.00 - 0.70 nm 95% CI) (Fig 3C). The reduction in distance parameter indicates that the ADP release rate of M493I-cHMM is modestly less sensitive to force than WT-cHMM, a possible contributing factor to asynchronous contraction in heterozygous disease myocardium. These effects on force production and ADP release may contribute to gain of function, which leads to septal hypertrophy and ultimately outflow restriction present in M493I patients.

### M493I enhances steady-state ATPase rates

The actin-activated steady-state ATPase rate was measured using a NADH-coupled ATPase assay, over a range of concentrations of porcine cardiac thin filaments (45). NADH absorbance decreases linearly over time with the slope indicating ATPase rate increasing with thin filament concentration (Fig. S3A-B). M493I myosins demonstrated higher ATPase rate than WT (Fig S3A-C). The *V_max_* of the per-head ATPase rate for WT-cHMM was 1.20 s^−1^ (1.09 - 1.77 s^−1^ 95% CI) while for M493I-cHMM it nearly doubled to 2.35 s^−1^ (1.51 - 2.51 s^−1^ 95% CI) without significantly altering the *K*_m_ (6.66 µM, 5.25 - 13.62 µM 95% CI for WT *vs.* 7.65 µM, 2.97 - 10.09 µM 95% CI for M493I). Thus, the slowed *in vitro* gliding filament motility is not caused by defective ATP cycling, as this rate is enhanced by the mutation (Fig S3C, Table 1).

### Absent actin, M493I enhances ATP turnover and reduces the equilibrium population of super-relaxed (SRX) myosin heads

Enhancement of *V_max_* for M493I-cHMM steady-state ATPase seems in conflict with the slowed ADP release observed from pyrene and optical trapping experiments, which should tend to slow cycling through the ATPase pathway. A possible explanation for this discrepancy would be a difference in the number of available catalytic myosin heads. Class II myosins from across animalia have a conserved off state, in which the two heads of myosin fold onto the proximal coiled-coil tail, forming stabilizing interactions with both the other head and the tail domain (Fig S1C) (31). In striated muscle myosins, this state is termed the interacting-head motif (IHM), and it has been correlated with a biochemically-defined off state termed the super-relaxed (SRX) state (32, 34–41, 59). In contrast, myosin heads in the alternative disordered-relaxed (DRX) state are available to interact with the thin filament.

A common method for measuring the SRX/DRX partition is through single-turnover studies using a fluorescent ATP analog, N-methylanthraniloyl adenosine 5’-triphosphate (mantATP). In this approach, the proportion of fast and slow release of mantADP, either with purified myosin constructs or in isolated myofibrils/myocytes is estimated by fitting the fluorescence change upon mantADP dissociation, as a proxy for nucleotide turnover, to a double exponential model (32, 34–36). For purified HMM, the presence of two-phase nucleotide release has been interpreted as indicating that the transition out of SRX is rate-limiting (38–41). Recent work from our group and others has found instead that nucleotide release occurs in a single phase, implying a rapid equilibrium between the SRX and DRX states (60, 61).

To assess the effect of the M493I mutation on the partitioning between the super-relaxed (SRX) and disordered relaxed (DRX) states, we performed single-nucleotide turnover experiments using mantATP (Fig. 4A). In this assay, purified cHMM was rapidly mixed with a slight excess (1.1-fold) of mantATP, aged for 10 seconds to allow nucleotide binding and hydrolysis, and then chased with saturating unlabeled ATP (1 mM). Tryptophan 508 in the active site was excited at 295 nm, resulting in fluorescence resonance energy transfer (FRET) excitation of the bound mant fluorophore. Upon nucleotide dissociation, fluorescence intensity decreased, and the decay in signal was fit to a single-exponential function corresponding to the basal rate of nucleotide release.

**Fig 4:**
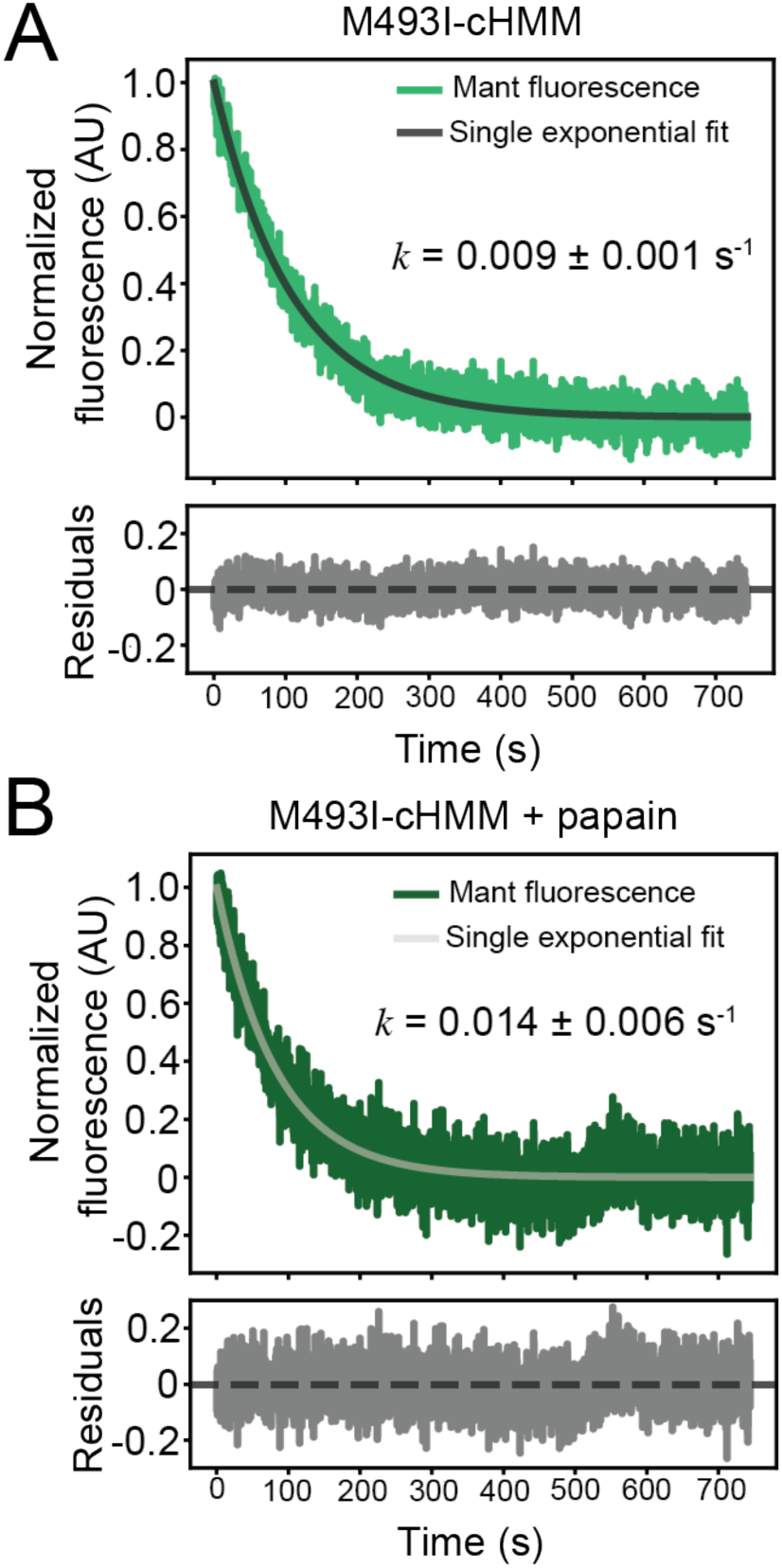
Turnover of MantATP for WT and M493I myosins. A) M493I-cHMM MantADP release with single exponential fit. B) MantADP release rate for M493I-cHMM after papain digestion, showing increased dissociation rate of release attributed to minimized (or lack of) population of SRX.

In experiments with soluble cardiac myosin, we found that correction for photobleaching eliminated the previously reported slower phase attributed to a slow SRX-to-DRX transition.

Instead, the presence of the S2 tail establishes a dynamic equilibrium between SRX and DRX that is faster than the rate of nucleotide turnover itself, as summarized by the following scheme:

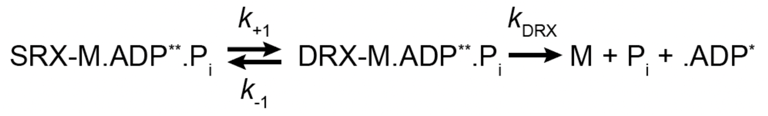

where M is myosin, ADP** is the high-fluorescence mantADP in the active site and ADP* is the low-fluorescence mantADP in solution (60). The equilibrium constant was determined according to:

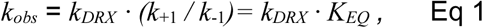

where *k_DRX_* is the elementary product release rate measured from cS1*, k*_+1_ and *k*_-1_ are the interconversion rates between SRX and DRX, and *K_EQ_ = k*_+1_ / *k*_-1_.

As determined previously, WT-cHMM releases nucleotide in a single phase at a rate of 0.0047 ± 0.0005 s^−1^ (60). WT-subfragment 1, either recombinantly expressed or produced by limited papain digestion which cleaves HMM into S1, S1, and single headed fragments (59), releases nucleotide at the significantly increased rate of 0.013 ± 0.001 s^−1^, from which we calculate an equilibrium constant *K*_SRX/DRX_ = 0.33 ± 0.05.

M493I-cHMM releases nucleotide with single-exponential kinetics, with a rate of 0.009 ± 0.001 s^−1^, approximately double that of WT-cHMM (Fig 4A). Upon papain digestion, the nucleotide release rate again increased to 0.014 ± 0.006 s^−1^, quite similar to the value for papain-digested WT-cHMM (Fig 4B, Fig S4A). Paired undigested M493I-cHMM control samples again displayed a similar rate constant to untreated samples (Fig S4B). Thus, we estimate that M493I-cHMM is in equilibrium between SRX and DRX at a rate substantially faster than the ATP turnover rate with an equilibrium constant *K*_SRX/DRX_ = 0.63 ± 0.04, nearly double that of WT-cHMM. The higher availability of DRX heads actively hydrolyzing ATP provides an explanation for the higher steady-state ATPase rate of the mutant myosin despite the slower ADP release.

### M493I enhances single-molecule actin rebinding rate

The increased availability of DRX heads in M493I-cHMM is expected to increase interactions with actin filaments. To assess this possibility, we performed single-molecule actomyosin interaction assays at saturating (2mM) ATP in our three-bead optical trap assay. In order to reliably measure the ON-rate of myosin to actin in the 3-bead assay, the technique was enhanced with a high-gain stage feedback system to maintain nanometer-scale precision in localizing the actin interaction zone (62) and careful optimization of each actin dumbbell for interaction with the myosin on the bead. Over the course of an acquisition of single actomyosin interactions, the actin filament position is centered and maintained relative to the cHMM (within the Brownian distribution of the optically trapped dumbbell), and the antibody-epitope adhesion scheme preserves the proximal S2 tail of the cHMM for forming SRX interactions (Fig 5A).

**Fig 5:**
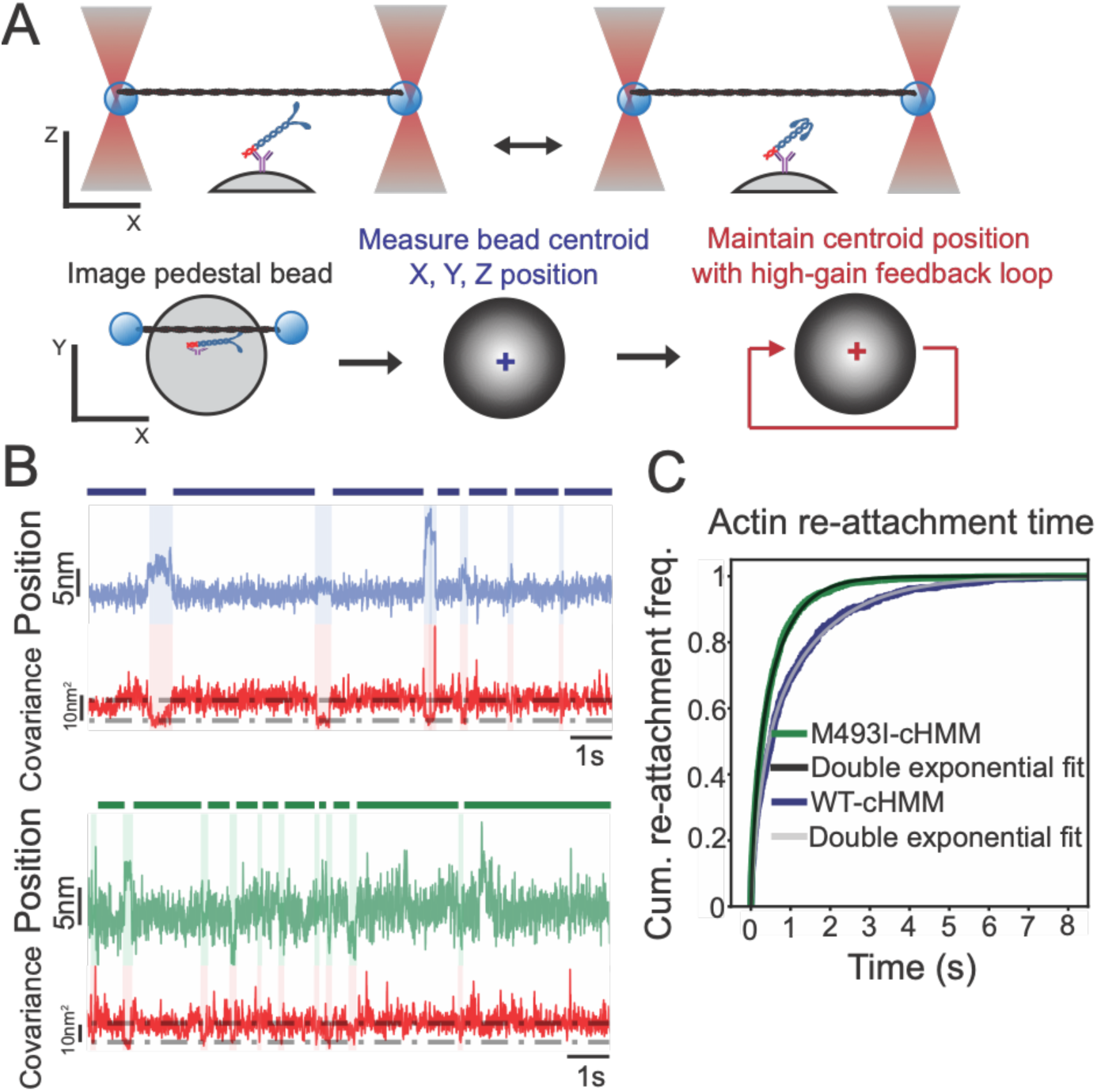
Actin reattachment rate measured by optical trap assays. A) Schematic of myosin immobilization that preserves S1-S2 interactions for SRX formation, and high-gain feedback loop to stabilize the actin-myosin interaction zone. B) Sample optical trap recordings showing faster re-attachment to actin for M493I myosin. C) Cumulative distributions of re-attachment times for M493I-cHMM and WT-cHMM showing faster re-attachment rate for M493I myosin relative to WT.

In direct comparison, traces of WT-cHMM interactions showed a significantly slower reattachment to actin filaments (blue lines, Fig. 5B) than those of M493I-cHMM (green lines, Fig. 5B). Analysis of reattachment times across multiple molecules revealed a double-exponential distribution of attachment rates. Both WT-cHMM and M493I-cHMM shared a minor kinetic component comprising ∼20% of the total amplitude, with similar rates. However, the dominant kinetic component, accounting for the majority of events, was nearly twice as fast in M493I-cHMM (1.62 s⁻¹, 95% CI: 1.45–1.74 s⁻¹) compared to WT-cHMM (0.83 s⁻¹, 95% CI: 0.70–0.90 s⁻¹, Fig. 5C). This suggests that the increased population of DRX heads in the M493I mutant enhances its ability to rapidly rebind actin.

## Discussion

Hypertrophic cardiomyopathy (HCM) is often described as a disease of hypercontractility, yet studies of individual purified myosin mutations have revealed mixed effects. Some mutations enhance motor activity, while others inhibit it. In this study, we characterized the mechanochemistry of a highly penetrant HCM mutation, M493I, and reconciled its seemingly contradictory kinetics. Although the mutation appears to reduce activity based on slower actin gliding motility, it ultimately leads to stronger and more sustained actin interactions: the mutant shows increased actin binding due to a two-fold disruption in super-relaxed (SRX) state formation, prolonged attachment times driven by a five-fold slowing of ADP release, and elevated force production by actomyosin cross-bridges. These perturbations, in the context of a heterozygous myocardium, are likely to result in asynchronous contraction, aberrant thin filament activation, and septal stiffening hypertrophy, as seen in patients.

### A relay helix point mutation disrupts the kinetic profile of cardiac myosin

The relay helices of myosins are tightly conserved across the human genome; the position corresponding to M493 is one of the most variable residues in *H. sapiens* myosin sequences, but appear as a methionine in all class II myosins. In the transport myosins (classes V and VI), this position bears a valine or isoleucine, as in M493I. These myosins release ADP slowly, as their rate-limiting step, a key factor in their high duty ratio and long actin attachment duration. In fact, the ADP release rate from AM·ADP of myosin V, at 11/s, is remarkably similar to the measured rate for M493I (63). M493I is also rate-limited by ADP release, rather than P_i_ release, which is the rate-limiting step for WT cardiac myosin (47). In myosin IB, the exemplary force-sensing myosin, a threonine occupies this position (64). These differences in relay helix characteristic, with corresponding changes to myosin activity, point to the important allosteric effects of the relay helix in communicating between the nucleotide active site and the myosin lever arm.

### Enhanced attachment duration leads to high loaded force production

The tight regulation of molecular interactions underpinning the cross bridge cycling that powers cardiac function implies that dysfunction in diseases such as HCM result from a range of effects of mutations on myosin function that may lead to hypercontractility. Here, the presence of long actin attachment durations from slowed ADP release may be related to significantly enhanced force production by M493I myosin under hindering load, possibly through the recruitment of the second myosin head. In the myocardium, this could lead to hypercontractility not only through increased force production by individual motors but also through the cooperative action of enhanced thin filament activation, inducing WT motors expressed from the other allele to contribute aberrantly high force. These effects may contribute to significant stiffening of the myocardium and imbalances in forces between neighboring myocytes, leading to asynchronous contraction, disorder, and interstitial fibrosis.

### Disruptions to the super-relaxed state enhance actin interactions

HCM mutations do not universally alter the SRX-DRX equilibrium of purified, soluble myosin. Previous work from our group has found, for instance, that the converter domain mutation R712L has no effect on SRX-DRX equilibrium, while E497D (also found in the relay helix) increases the equilibrium constant of the SRX-DRX equilibrium by approximately 50% (60). Thus, the hypothesis that an increase in available DRX heads contributes to HCM is not universally true for soluble myosin. The SRX-DRX transition may be different in a native thick filament, so additional experiments need to be performed in situ. The mutation M493I approximately doubles the equilibrium constant of the SRX/DRX transition, consistent with the head availability model of HCM. This mutation also doubles *V_max_* of actin-activated ATP hydrolysis measured in bulk kinetics, as expected with more available motor domains, and doubles the actin re-attachment rate measured in single-molecule optical trap assays. With the compounding impacts of disruption to the SRX “off” state on two-headed myosins with higher duty ratio than WT, small effects on mechanical and kinetic transitions are amplified to create significant composite changes to myosin function.

## Materials and Methods

### Protein purification

Heavy meromyosin (HMM) of human β-cardiac myosin (MYH7) was expressed in mouse C2C12 myoblasts and purified according to previously established methods (16, 25, 46, 60, 65). Briefly, the cHMM cDNA was cloned into the pShuttle-IRES-hrGFP-1 vector (Agilent Tech., Santa Clara, CA). The AdcHMM-Flag virus was prepared and amplified for expression of cHMM protein in C2C12 cells. For the cHMM2.0 construct, the sequence of the epitope (AEKHRADLSRE) was introduced into the coiled-coil S2 domain of β-cHMM, followed by two additional heptads of cardiac S2 sequence and a FLAG tag at the C-terminus (42). Mutant adenovirus constructs were synthesized by Genewiz (South Plainfield, NJ). The virus was expanded by infection of a large number of plates of confluent Ad293 cells at multiplicity of infection (MOI) of 3–5. The virus was harvested from the cells and purified by CsCl density sedimentation yielding final virus titers of 10^10^–10^11^ plaque forming units per mL (pfu⋅mL^−1^). Confluent C2C12 myoblasts were infected with replication defective recombinant adenovirus (AdcHMM-Flag) at 2.7 × 10^8^ pfu⋅mL^−1^ in fusion medium (89% DMEM, 10% horse serum, 1% FBS). Expression of recombinant cHMM was monitored by accumulation of co-expressed GFP fluorescence in infected cells. Myocyte differentiation and GFP accumulation were monitored for 216–264 hr after which the cells were harvested. Cells were chilled, media removed, and the cell layer was rinsed with cold PBS. The cell layer was scraped into Triton extraction buffer: 100 mM NaCl, 0.5% Triton X-100, 10 mM Imidazole pH 7.0, 1 mM DTT, 5 mM MgATP, and protease inhibitor cocktail (Sigma, St. Louis, MO). The cell suspension was collected in an ice-cold Dounce homogenizer and lysed with 15 strokes of the tight pestle. The cell debris in the whole cell lysate was pelleted by centrifugation at 17,000 x g for 15 min at 4°C. The Triton soluble extract was fractionated by ammonium sulfate precipitation using sequential steps of 0–30% and 30–60% saturation. The cHMM precipitates between 30–60% saturation of ammonium sulfate. The recovered pellet was dissolved in and dialyzed against 10 mM Imidazole, 150 mM NaCl, pH 7.4 for affinity purification of the FLAG-tagged cHMM on M2 mAb-Sepharose beads (Sigma). Bound cHMM was eluted with 0.1 mg/mL FLAG peptide (Sigma). Protein was concentrated and buffer exchanged on Amicon Ultracel-10K centrifugal filters (Millipore; Darmstadt, Germany), dialyzed exhaustively into 10 mM MOPS, 100 mM KCl, 1 mM DTT before a final centrifugation at 300,000 x g for 10 min at 4°C. Aliquots were drop frozen in liquid nitrogen and stored in vapor phase at –147°C. Purified WT human β-cHMM and M493I HCM variants were routinely analyzed by SDS-PAGE.

Actin was purified from rabbit skeletal muscle (66). Native porcine cardiac thin filaments (TFs) were prepared according to the procedure of Spiess et al., 1999 (67) as modified by Matsumoto et al., 2004 (68).

### Protein gels

Proteins were prepared in 1X Laemmli buffer with 0.5% DTT and run on NuPAGE Bis-Tris acrylamide gels 4-12% (Invitrogen), followed by Coomassie staining and destaining according to standard procedures. Gels were imaged on a Licor Odyssey gel imaging dock; all gel images are unedited and uncropped except to exclude irrelevant lanes.

### In vitro gliding assays

Measurement of in vitro motility of human β-cHMM2.0 was done as previously described for skeletal muscle myosin (16, 25, 69, 70). Nitrocellulose-coated glass coverslips were incubated with 0.15 mg/mL of the mAb 10F12.3, followed by blocking the surface with 1% BSA. β-cHMM proteins were diluted in motility buffer (MB) (25 mM imidazole, pH 7.8, 25 mM KCl, 4 mM MgCl2, 1 mM MgATP, 1 mM DTT) supplemented with 1% BSA (MB/BSA) to a final concentration of 10 μg/mL. The antibody-coated coverslips were incubated with β-cHMM2.0 for ∼2 h in a humidified chamber at 4 °C. The coverslips were washed with MB/BSA, followed by actin blocking with 1 μM F-actin, and washes with motility buffer, then transferred to a 15-μL drop of 2 nM rhodamine-phalloidin–labeled actin in a modified motility buffer (with 7.6 mM MgATP, 50 mM DTT, 0.5% methyl cellulose, 0.1 mg/mL glucose oxidase, 0.018 mg/mL catalase, 2.3 mg/mL glucose) in a small parafilm ring fixed on an alumina slide with vacuum grease. The chamber was observed with a temperature-controlled stage and objective set at 32°C on an upright microscope with an image-intensified charge-coupled device camera capturing data to an acquisition computer at 30 Hz. Movement of actin filaments from 500 to 1,000 frames of continuous imaging was analyzed with semiautomated filament tracking programs as previously described (69). The trajectory of every filament with a lifetime of at least 10 frames was determined; the instantaneous velocity of the filament moving along the trajectory, the filament length, the distance of continuous motion and the duration of pauses were tabulated. A weighted probability of the actin filament velocity for hundreds of events was fit to a Gaussian distribution and reported as a mean velocity and SD for each experimental condition. For presentation purposes, background was subtracted using a rolling ball of 1 pixel radius with sliding paraboloid shape, followed by Gaussian smoothing of size 3×3 pixels, both in ImageJ. Color-coded time projections were produced with the ImageJ plugin ColorCodingFrames.ijm (https://github.com/hansenjn/ColorStackByTimeAndProject).

### Three-bead optical trap assay

Optical trap assays were performed at room temperature (20° C) in flow cell chambers constructed of a microscope slide (75 x 25 mm) with coverslip (40 x 25 mm) adhered by double-sided tape. The coverslip was coated with 0.1% nitrocellulose (EMS) in amyl acetate (EMS) including dilute 2.5 μm silica pedestal beads (Polysciences). All reagents were in Assay Buffer (AB) which contained 60 mM MOPS, 25 mM KCL, 1 mM DTT, 1 mM MgCl_2_, and 1 mM EGTA unless otherwise specified. Antibodies to the HMM 10F12.3 epitope tag were diluted to 0.03 mg/mL and incubated for 60 s to adhere to the coverslip surface. The chamber was then blocked 2x for 3 minutes each in 1 mg/mL BSA, then incubated in cHMM (WT or M493I) diluted to 0.05 - 0.1 μg/mL in myosin buffer (AB except with 300 mM KCl) for 3 minutes, then blocked 2x for 2 minutes each in 1 mg/mL BSA. The experimental solution was added to the flow cell containing 0.2 nM actin filaments containing 10-15% biotinylated actin and 85-90% rabbit skeletal muscle actin, which was stabilized by equimolar rhodamine-labeled phalloidin, as well as an oxygen scavenging system plus glucose (100X GOC was prepared by dissolving 7.5 mg of glucose oxidase (Sigma-Aldrich G2133) in 200 uL of 10mM HEPES pH 7.4, and adding 60 uL of bovine liver catalase (Sigma-Aldrich C3155), spinning at 17,900xg for 5 minutes, and filtering through 0.22 um filter unit (EMD Millipore SLGV004SL)., and varying concentrations of MgATP. Finally, 750 nm polystyrene beads, prepared by incubating 0.4 ng of beads with 10 μL of 10 mg/mL neutravidin solution in water overnight at 4° C with rotation before washing 2x with AB+MgATP were diluted 1:100 in AB+MgATP. 3 μL of the bead suspension was added to the chamber before it was sealed with vacuum grease. Chambers were imaged for <60 minutes after sealing. Concentrations of antibody and myosin were optimized such that one of each 5-10 locations tested showed actomyosin interactions. Chambers constructed with no antibody in the first flow step, but following all other steps the same, showed no actomyosin interactions.

Experiments were performed on a dual-beam optical trap setup constructed on a Nikon Eclipse TE2000 microscope. A 1064-nm laser was split into two by polarization, with each beam passing through a 1-D electro-optical deflector, each of which can deflect the beam based on voltage. The position of one trap was also varied by rotating a motorized mirror conjugate to the back focal plane of the objective. The beams were focused at the focal plane with a Nikon Plan Apo 60x water immersion objective (1.2 NA), and the traps were projected by a Nikon HNA oil condenser (1.4 NA) onto two quadrant photodiodes (JQ-50P, Electro Optical Components Inc.) for position and force detection. Data acquisition, feedback, and beam position were controlled with custom-built virtual instruments programmed on a LabVIEW multi-function I/O device with a built-in FPGA (PXI-7851); data were acquired at 250 kHz. Beads were caught by moving the microscope stage to steer them into the traps. Stiffness of the traps were 0.07-0.1 pN/nm, as calculated via the power spectrum of the beads’ fluctuations. An actin filament 5-10 μm long was then suspended between the two beads, and the position of one trap was moved away from the other with the motorized mirror conjugate until 4.5-5.5 pN of pre-tension was applied to the actin filament. Pedestal beads were tested by pressing the actin filament against the pedestal to detect bead variance changes. Once actomyosin interactions were detected, a piezo-electric stage (Mad City Labs) was used to optimize the interaction zone, by visualizing both number (for high [ATP]) and bead displacement (for low [ATP]) of attachment events.

A stage feedback system imaged the pedestal bead with a monochrome camera (Thorlabs) as described previously (62). The gain of the feedback loop was set to maintain <3 nm error in X and Y positions, and <10 nm error in Z position. Upon engagement of the feedback loop, data were acquired without adjustment to bead positions.

Isometric feedback experiments were conducted using a digital feedback loop and the EODs to steer the beam position (57). A feedback loop held the position of the pointed-end associated bead (referred to as the transducer) constant by modulating the position of the trap holding barbed-end associated bead, termined the motor trap. Since the actin filament is between the two beads inside the feedback loop, its position was maintained continuously, allowing the myosin to develop isometric force during its interaction with actin. The response time of the feedback loop during myosin interactions was ∼10 ms. The average force during an interaction was calculated by averaging the force on the motor bead starting 2 ms after detected attachment through 2 ms before detachment. The baseline force on the motor bead 4 ms after detachment was subtracted from this average force.

### Optical trap data analysis

Optical trap data were analyzed using python scripts available at GitHub.com/bobcail. First, the running covariance of the beads from a 15 s recording was calculated over an 8 ms sliding window and fit by a double Gaussian distribution, with high-covariance values corresponding to unbound actin and low-covariance values corresponding to actomyosin interaction events. Events were selected by finding where the covariance dropped below the high peak, then below the lower peak, then rose above the high peak again. Events shorter than 16 ms (the dead time of the instrument) were excluded. The duration of each event was calculated from these trigger points; event durations were fit by single exponential functions using MEMLET (71). Substep sizes were calculated from 1 μM ATP data by averaging 1 ms of the bead position at three points: 1) 4 ms prior to attachment, 2) 2 ms after attachment, and 3) 4 ms before detachment. Substep 1 was calculated as 2-1, and substep 2 was calculated as 3 - 2. Ensemble averaging was performed by aligning all events to the start (forward average) or stop (reverse average), and events were extended to the longest event time by averaging 4 ms of data, located 1 ms from the covariance transition point. Events were averaged with each event contributing equal weight to the ensemble. Ensembles were fit by single exponential functions by nonlinear least squares fitting in python. All plots were generated in python.

### Stopped-flow experiments

Stopped flow experiments were conducted in an SX20 Stopped Flow Spectrometer (Applied Photophysics), with fluorescence excitation provided by a 100-W Hg lamp with a monochromator. Data were acquired and analyzed using Pro Data-SX software. Pyrene-actin was produced following previously published protocols (45). Pyrene-actin for ADP release and ATP-induced actomyosin dissociation and was excited at 365 nm, and the fluorescence emission peak was detected through a 400 nm long-pass filter. For ADP release, 1 μM M493I or 200 nM WT myosin heads (incubated with 0.01 U/mL apyrase on ice for 30 minutes) was mixed with equimolar pyrene-actin with 280 μM ADP for 10 minutes at RT in KMg25 buffer (60 mM 3-(N-morpholino)propanesulfonic acid, pH 7.0, 1 mM MgCl_2_, 1 mM EGTA, and 1 mM DTT), then mixed with 4 mM ATP in KMg25 and fluorescence intensity was observed for 1 second. Final concentrations in cuvette: 0.5 μM heads, 0.5 μM pyrene-actin, 140 μM ADP, 2 mM ATP for M493I, 0.1 μM heads, 0.1 μM pyrene-actin, 140 μM ADP, 2 mM ATP for WT. For ATP binding, 1 μM myosin heads (incubated with 0.01 U/mL apyrase on ice for 30 minutes) were mixed with 1 μM pyrene-actin for 10 minutes at RT in KMg25 buffer (60 mM MOPS pH 7, 25 mM KCl, 1 mM EGTA, 1 mM MgCl2, 1 mM DTT), then mixed with 0-1200 μM ATP in KMg25 and fluorescence intensity was observed for 1 second. Final concentrations in cuvette: 0.5 μM heads, 0.5 μM pyrene-actin, 0-600 μM ATP. For actin-activated phosphate release, fluorescently labeled mutant phosphate binding protein (MDCC-labeled PiBiP; (7-diethylamino-3-((((2-maleimidyl)ethyl)amino)carbonyl) coumarin)-labeled phosphate binding protein) was excited at 430 nm, and fluorescence was detected with a 440 nm long-pass filter. The instrument was pre-cleaned with a phosphate mop consisting of 0.3 mM 7-methylguanosine and 0.1 U/mL bacterial nucleoside phosphorylase in phosphate release buffer (0.11 mM CaCl_2_, 10 mM KCL, 2 mM MgCl_2_, 10 mM MOPS pH 7.2). Phosphate release was measured in phosphate release buffer with 5 μM MDCC-labeled PiBiP, 0.1mM 7-methylguanosine and 0.01 U/mL bacterial nucleoside phosphorylase; 4 μM myosin heads were mixed with 4.1 μM ATP in the pre-incubation loop for 10 seconds, followed by mixing with 60 μM TFs, and fluorescence measured for 21 seconds on a split time base (1000 points in first second, 1000 points for 20 seconds). Final concentrations in cuvette: 1 μM myosin heads, 1.025 μM ATP, 30 μM TFs.

For single-nucleotide turnover, 200 nM myosin heads were incubated with 200 nM MantATP in the pre-incubation loop for 10 seconds, followed by mixing with 2 mM unlabeled ATP. Mant-ATP was excited by FRET from tryptophan 508 with 295 nm excitation, and fluorescence was collected with a 400 nm long-pass filter. Final concentration in cuvette: 50 nM myosin heads, 50 nM MantATP, 1 mM unlabeled ATP. For proteolytic fragment formation, 0.5 mg/mL HMM (apyrase treated at 0.1 U/mL on ice for 30 minutes before use) was incubated in a total of 60 μL KMg25 with 5 mM cysteine and 18.1 μM papain (diluted from 1.09 mM stock, Sigma-Aldrich p3125) for 5 minutes for WT or 3 minutes for M493I at room temperature. The reaction was quenched with addition of 25 μM E-64 protease inhibitor (Cayman Chemical), and 10 μL was reserved for SDS-PAGE analysis; in parallel, a sample was prepared identically but without the addition of papain, to test samples and the effect of incubation times and the E-64. Samples were incubated on ice until single-nucleotide turnover, which was performed identically as for untreated samples. Stopped-flow data were fitted by single exponentials and rates were fitted where appropriate by the Michaelis-Menten equation using custom Python scripts.

### Actin-activated ATP hydrolysis

Steady state ATPase experiments were measured spectrophotometrically (Agilent) using to standard protocols. Briefly, an assay buffer containing 0.11 mM CaCl_2_, 10 mM KCL, 2 mM MgCl_2_, 10 mM MOPS pH 7.2, 2mM MgATP, 0.2 mM NADH, 20 U/mL lactate dehydrogenase, 100 U/mL pyruvate kinase, and 0.5 mM phospho(enol)pyruvate was incubated with 100 nM myosin heads 0-60 μM TFs (all final concentration) and the absorbance was measured at 340 nM over 110 seconds. The slope of the fluorescence change was fitted by a straight line, the slope giving the rate of decrease in NADH concentration (extinction coefficient of 6220 M^−1^cm^−1^) which corresponds to the rate of ATP hydrolysis. Per-head ATPase rates were fitted to the Michaelis-Menten equation with custom python scripts.

### Statistical tests

Purified WT and M493I cHMM preparations represent 3 biological replicates. All measurements include 3 technical replicates. Measured values are reported as mean +/− S.D. unless otherwise stated. Fitted values are reported as value with 95% C.I. as determined by bootstrapping for 1000 iterations.

## Author Contributions

Experiments were conceptualized by E.M.O., Y.E.G., R.C.C., and D.A.W. Proteins were purified by D.A.W. and B.B. Motility assays were performed and analyzed by B.B. and D.A.W. Transient and steady-state kinetics were performed by R.C.C. and F.A.B.C. Optical trap experiments were performed by R.C.C. All data were analyzed by R.C.C. except motility assays. Figures were prepared by R.C.C. Manuscript was written by R.C.C. Y.E.G. and E.M.O. All authors contributed to manuscript editing.

## Data availability

Representative traces of kinetics and optical trap acquisitions, representative micrographs of fluorescence microscopy data, and unedited SDS-PAGE/Coomassie gels are presented in this manuscript; all fits and statistics are reported, and all data are included. Raw data are available upon request from R.C.C. and E.M.O. Code for analysis is available at GitHub.com/bobcail.

## Acknowledgements

This work was supported by the Center for Engineering MechanoBiology NSF Science and Technology Center, CMMI: 15-48571 to Y.E.G. and E.M.O, National Institutes of Health grants 5R01HL157997-04 to D.A.W., Y.E.G. and E.M.O., R35GM118139 to Y.E.G., R37GM057247 to E.M.O., and 5T32AR053461 to R.C.C. We thank Daniel Safer for technical assistance.

## Notes

### Competing Interest Statement

The authors have declared no competing interest.

### Summary of Updates

Updated graph to better represent force dependent detachment.

## References

1. B. J. Maron, et al., Prevalence of Hypertrophic Cardiomyopathy in a General Population of Young Adults. Circulation 92, 785–789 (1995).

2. B. J. Maron, M. S. Maron, Hypertrophic cardiomyopathy. The Lancet 381, 242–255 (2013).

3. A. A. Young, C. M. Kramer, V. A. Ferrari, L. Axel, N. Reichek, Three-dimensional left ventricular deformation in hypertrophic cardiomyopathy. Circulation 90, 854–867 (1994).

4. A. M. Varnava, P. M. Elliott, N. Mahon, M. J. Davies, W. J. McKenna, Relation between myocyte disarray and outcome in hypertrophic cardiomyopathy. The American Journal of Cardiology 88, 275–279 (2001).

5. G. Moravsky, et al., Myocardial Fibrosis in Hypertrophic Cardiomyopathy: Accurate Reflection of Histopathological Findings by CMR. JACC: Cardiovascular Imaging (2013). 10.1016/j.jcmg.2012.09.018.

6. C. Y. Ho, et al., Myocardial Fibrosis as an Early Manifestation of Hypertrophic Cardiomyopathy. New England Journal of Medicine 363, 552–563 (2010).

7. E. Braunwald, C. T. Lambrew, S. D. Rockoff, J. Ross, A. G. Morrow, Idiopathic Hypertrophic Subaortic Stenosis: I. A Description of the Disease Based Upon an Analysis of 64 Patients. Circulation 29, IV–3 (1964).

8. G. M. Hecht, H. G. Klues, W. C. Roberts, B. J. Maron, Coexistence of sudden cardiac death and end-stage heart failure in familial hypertrophic cardiomyopathy. J Am Coll Cardiol 22, 489–497 (1993).

9. B. J. Maron, M. S. Maron, C. Semsarian, Genetics of Hypertrophic Cardiomyopathy After 20 Years: Clinical Perspectives. Journal of the American College of Cardiology (2012). 10.1016/j.jacc.2012.02.068.

10. S. P. Harris, R. G. Lyons, K. L. Bezold, In the Thick of It: HCM-Causing Mutations in Myosin Binding Proteins of the Thick Filament. Circulation research 108, 751 (2011).

11. S. L. Van Driest, et al., Comprehensive analysis of the beta-myosin heavy chain gene in 389 unrelated patients with hypertrophic cardiomyopathy. J Am Coll Cardiol 44, 602–610 (2004).

12. C. Y. Ho, et al., Echocardiographic Strain Imaging to Assess Early and Late Consequences of Sarcomere Mutations in Hypertrophic Cardiomyopathy. Circulation: Cardiovascular Genetics 2, 314–321 (2009).

13. C. Y. Ho, et al., Assessment of Diastolic Function With Doppler Tissue Imaging to Predict Genotype in Preclinical Hypertrophic Cardiomyopathy. Circulation 105, 2992–2997 (2002).

14. J. A. Spudich, Three perspectives on the molecular basis of hypercontractility caused by hypertrophic cardiomyopathy mutations. Pflugers Arch - Eur J Physiol 471, 701–717 (2019).

15. E. R. Witjas-Paalberends, et al., Mutations in MYH7 reduce the force generating capacity of sarcomeres in human familial hypertrophic cardiomyopathy. Cardiovascular Research 99, 432–441 (2013).

16. A. Snoberger, et al., Myosin with hypertrophic cardiac mutation R712L has a decreased working stroke which is rescued by omecamtiv mecarbil. eLife 10, e63691 (2021).

17. R. F. Sommese, et al., Molecular consequences of the R453C hypertrophic cardiomyopathy mutation on human β-cardiac myosin motor function. Proceedings of the National Academy of Sciences 110, 12607–12612 (2013).

18. C. D. Vera, et al., Myosin motor domains carrying mutations implicated in early or late onset hypertrophic cardiomyopathy have similar properties. Journal of Biological Chemistry 294, 17451–17462 (2019).

19. S. Nag, et al., Contractility parameters of human β-cardiac myosin with the hypertrophic cardiomyopathy mutation R403Q show loss of motor function. Science Advances (2015). 10.1126/sciadv.1500511.

20. T. G. M. Oliveira, et al., A Variant Detection Pipeline for Inherited Cardiomyopathy-Associated Genes Using Next-Generation Sequencing. J Mol Diagn 17, 420–430 (2015).

21. J. D. C. Marsiglia, et al., Screening of MYH7, MYBPC3, and TNNT2 genes in Brazilian patients with hypertrophic cardiomyopathy. American Heart Journal 166, 775–782 (2013).

22. V. J. Planelles-Herrero, J. J. Hartman, J. Robert-Paganin, F. I. Malik, A. Houdusse, Mechanistic and structural basis for activation of cardiac myosin force production by omecamtiv mecarbil. Nat Commun 8, 190 (2017).

23. N. Nandwani, et al., Hypertrophic cardiomyopathy mutations Y115H and E497D disrupt the folded-back state of human β-cardiac myosin allosterically. [Preprint] (2024). Available at: https://www.biorxiv.org/content/10.1101/2024.02.29.582851v1

24. M. H. Doran, et al., Conformational changes linked to ADP release from human cardiac myosin bound to actin-tropomyosin | Journal of General Physiology | Rockefeller University Press. 10.1085/jgp.202213267.

25. D. A. Winkelmann, E. Forgacs, M. T. Miller, A. M. Stock, Structural basis for drug-induced allosteric changes to human β-cardiac myosin motor activity. Nat Commun 6, 7974 (2015).

26. B. M. Palmer, et al., Elevated rates of force development and MgATP binding in F764L and S532P myosin mutations causing dilated cardiomyopathy. Journal of Molecular and Cellular Cardiology 57, 23–31 (2013).

27. R. V. Agafonov, et al., Structural dynamics of the myosin relay helix by time-resolved EPR and FRET. Proceedings of the National Academy of Sciences 106, 21625–21630 (2009).

28. W. M. Shih, J. A. Spudich, The Myosin Relay Helix to Converter Interface Remains Intact throughout the Actomyosin ATPase Cycle *. Journal of Biological Chemistry 276, 19491–19494 (2001).

29. L. M. Coluccio, Myosins: A Superfamily of Molecular Motors (Springer Netherlands, 2008).

30. H. L. Sweeney, A. Houdusse, Structural and Functional Insights into the Myosin Motor Mechanism. Annual Review of Biophysics 39, 539–557 (2010).

31. K. H. Lee, et al., Interacting-heads motif has been conserved as a mechanism of myosin II inhibition since before the origin of animals. Proceedings of the National Academy of Sciences 115, E1991–E2000 (2018).

32. H. S. Jung, S. Komatsu, M. Ikebe, R. Craig, Head–Head and Head–Tail Interaction: A General Mechanism for Switching Off Myosin II Activity in Cells. MBoC 19, 3234–3242 (2008).

33. K. M. Trybus, T. W. Huiatt, S. Lowey, A bent monomeric conformation of myosin from smooth muscle. Proceedings of the National Academy of Sciences 79, 6151–6155 (1982).

34. M. A. Stewart, K. Franks-Skiba, S. Chen, R. Cooke, Myosin ATP turnover rate is a mechanism involved in thermogenesis in resting skeletal muscle fibers. Proceedings of the National Academy of Sciences 107, 430–435 (2010).

35. R. L. Anderson, et al., Deciphering the super relaxed state of human β-cardiac myosin and the mode of action of mavacamten from myosin molecules to muscle fibers. Proceedings of the National Academy of Sciences 115, E8143–E8152 (2018).

36. P. Hooijman, M. A. Stewart, R. Cooke, A New State of Cardiac Myosin with Very Slow ATP Turnover: A Potential Cardioprotective Mechanism in the Heart. Biophysical Journal 100, 1969–1976 (2011).

37. A. Grinzato, et al., Cryo-EM structure of the folded-back state of human β-cardiac myosin. Nat Commun 14, 3166 (2023).

38. J. A. Spudich, N. Nandwani, J. Robert-Paganin, A. Houdusse, K. M. Ruppel, Reassessing the unifying hypothesis for hypercontractility caused by myosin mutations in hypertrophic cardiomyopathy. The EMBO Journal 43, 4139–4155 (2024).

39. J. A. Spudich, The myosin mesa and a possible unifying hypothesis for the molecular basis of human hypertrophic cardiomyopathy. Biochemical Society Transactions 43, 64–72 (2015).

40. J. A. Spudich, Hypertrophic and Dilated Cardiomyopathy: Four Decades of Basic Research on Muscle Lead to Potential Therapeutic Approaches to These Devastating Genetic Diseases. Biophysical Journal 106, 1236–1249 (2014).

41. C. N. Toepfer, et al., Myosin Sequestration Regulates Sarcomere Function, Cardiomyocyte Energetics, and Metabolism, Informing the Pathogenesis of Hypertrophic Cardiomyopathy. Circulation 141, 828–842 (2020).

42. D. A. Winkelmann, S. Lowey, Probing myosin head structure with monoclonal antibodies. Journal of Molecular Biology 188, 595–612 (1986).

43. D. E. Harris, D. M. Warshaw, Slowing of velocity during isotonic shortening in single isolated smooth muscle cells. Evidence for an internal load. Journal of General Physiology 96, 581–601 (1990).

44. D. E. Harris, S. S. Work, R. K. Wright, N. R. Alpert, D. M. Warshaw, Smooth, cardiac and skeletal muscle myosin force and motion generation assessed by cross-bridge mechanical interactions in vitro. J Muscle Res Cell Motil 15, 11–19 (1994).

45. E. M. De La Cruz, E. Michael Ostap, “Chapter 6 Kinetic and Equilibrium Analysis of the Myosin ATPase” in Methods in Enzymology, Biothermodynamics, Part A., (Academic Press, 2009), pp. 157–192.

46. M. Brune, J. L. Hunter, J. E. T. Corrie, M. R. Webb, Direct, real-time measurement of rapid inorganic phosphate release using a novel fluorescent probe and its application to actomyosin subfragment 1 ATPase. Biochemistry 33, 8262–8271 (1994).

47. Y. Liu, H. D. White, B. Belknap, D. A. Winkelmann, E. Forgacs, Omecamtiv Mecarbil Modulates the Kinetic and Motile Properties of Porcine β-Cardiac Myosin. Biochemistry 54, 1963–1975 (2015).

48. J. A. Rohde, D. D. Thomas, J. M. Muretta, Heart failure drug changes the mechanoenzymology of the cardiac myosin powerstroke. Proceedings of the National Academy of Sciences 114, E1796–E1804 (2017).

49. A. D. Mehta, J. T. Finer, J. A. Spudich, Detection of single-molecule interactions using correlated thermal diffusion. Proceedings of the National Academy of Sciences 94, 7927–7931 (1997).

50. J. M. Laakso, J. H. Lewis, H. Shuman, E. M. Ostap, Myosin I Can Act As a Molecular Force Sensor. Science 321, 133–136 (2008).

51. C. Veigel, et al., The motor protein myosin-I produces its working stroke in two steps. Nature 398, 530–533 (1999).

52. C. Chen, et al., Kinetic Schemes for Post-Synchronized Single Molecule Dynamics. Biophysical Journal 102, L23–L25 (2012).

53. M. J. Greenberg, H. Shuman, E. M. Ostap, Inherent Force-Dependent Properties of β-Cardiac Myosin Contribute to the Force-Velocity Relationship of Cardiac Muscle. Biophysical Journal 107, L41–L44 (2014).

54. C. Veigel, J. E. Molloy, S. Schmitz, J. Kendrick-Jones, Load-dependent kinetics of force production by smooth muscle myosin measured with optical tweezers. Nature Cell Biology 5, 980–986 (2003).

55. M. Caremani, L. Melli, M. Dolfi, V. Lombardi, M. Linari, Force and number of myosin motors during muscle shortening and the coupling with the release of the ATP hydrolysis products. The Journal of Physiology 593, 3313–3332 (2015).

56. W. O. Fenn, B. S. Marsh, Muscular force at different speeds of shortening. The Journal of Physiology 85, 277 (1935).

57. Y. Takagi, E. E. Homsher, Y. E. Goldman, H. Shuman, Force Generation in Single Conventional Actomyosin Complexes under High Dynamic Load. Biophysical Journal 90, 1295–1307 (2006).

58. Bell, G.I, Models for the Specific Adhesion of Cells to Cells | Science. Available at: https://www.science.org/doi/abs/10.1126/science.347575S

59. S. Duno-Miranda, et al., Tail length and E525K dilated cardiomyopathy mutant alter human β-cardiac myosin super-relaxed state. Journal of General Physiology 156, e202313522 (2024).

60. R. C. Cail, F. A. Baez-Cruz, D. A. Winkelmann, Y. E. Goldman, E. M. Ostap, Dynamics of β-cardiac myosin between the super-relaxed and disordered-relaxed states. Journal of Biological Chemistry 108412 (2025). 10.1016/j.jbc.2025.108412.

61. Mohran, et al., The biochemically defined super relaxed state of myosin—A paradox. Journal of Biological Chemistry 300 (2024).

62. M. Capitanio, R. Cicchi, F. S. Pavone, Position control and optical manipulation for nanotechnology applications. Eur. Phys. J. B 46, 1–8 (2005).

63. E. M. De La Cruz, A. L. Wells, S. S. Rosenfeld, E. M. Ostap, H. L. Sweeney, The kinetic mechanism of myosin V. Proceedings of the National Academy of Sciences 96, 13726–13731 (1999).

64. H. Shuman, et al., A vertebrate myosin-I structure reveals unique insights into myosin mechanochemical tuning. Proceedings of the National Academy of Sciences 111, 2116–2121 (2014).

65. M. S. Woody, et al., Positive cardiac inotrope omecamtiv mecarbil activates muscle despite suppressing the myosin working stroke. Nat Commun 9, 3838 (2018).

66. J. A. Spudich, S. Watt, The Regulation of Rabbit Skeletal Muscle Contraction. Journal of Biological Chemistry 246, 4866–4871 (1971).

67. M. Spiess, et al., Isolation, Electron Microscopic Imaging, and 3-D Visualization of Native Cardiac Thin Myofilaments. Journal of Structural Biology 126, 98–104 (1999).

68. F. Matsumoto, et al., Conformational Changes of Troponin C Within the Thin Filaments Detected by Neutron Scattering. Journal of Molecular Biology 342, 1209–1221 (2004).

69. B. Barua, D. A. Winkelmann, H. D. White, S. E. Hitchcock-DeGregori, Regulation of actin-myosin interaction by conserved periodic sites of tropomyosin. Proceedings of the National Academy of Sciences 109, 18425–18430 (2012).

70. L. Bourdieu, Spiral Defects in Motility Assays: A Measure of Motor Protein Force. Phys. Rev. Lett. 75, 176–179 (1995).

71. M. S. Woody, J. H. Lewis, M. J. Greenberg, Y. E. Goldman, E. M. Ostap, MEMLET: An Easy-to-Use Tool for Data Fitting and Model Comparison Using Maximum-Likelihood Estimation. Biophysical Journal 111, 273–282 (2016).

