## Supplementary material for "A myosin hypertrophic cardiomyopathy mutation disrupts the super-relaxed state and boosts contractility by enhanced actin attachment": Supp Figures

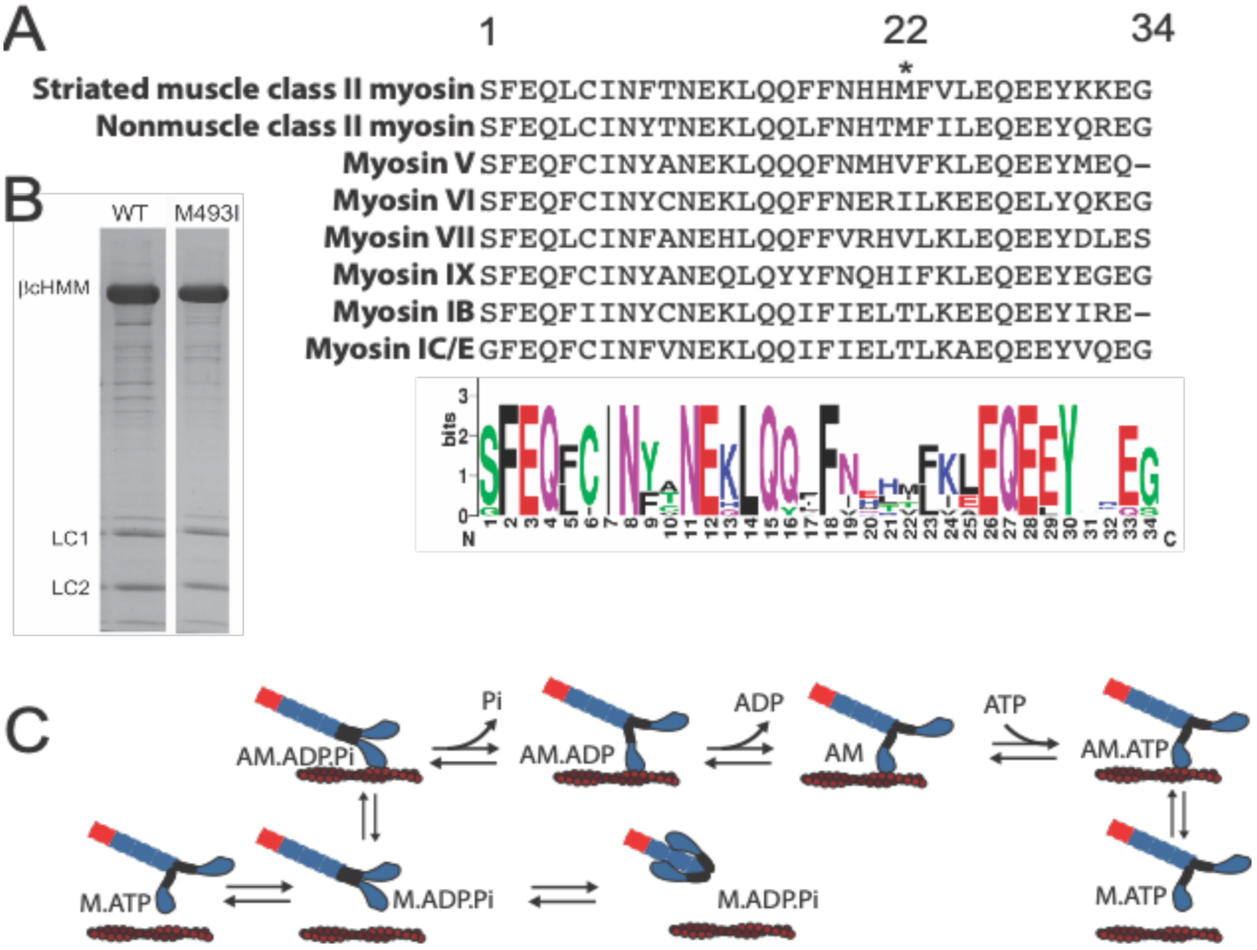


Fig S1: WT and M493I myosins. A) Comparison of relay helix sequences across many *H. sapiens* myosin paralogs. M493, at position 22 in the relay helix, is conserved in class II myosins but variable across paralogs. B) SDS-PAGE gels of purified WT- and M493I-cHMMs expressed recombinantly in mouse C2C12 myoblasts and purified. C) ATPase scheme of class II myosins, which includes the SRX-DRX regulatory transition for M.ADP.Pi.

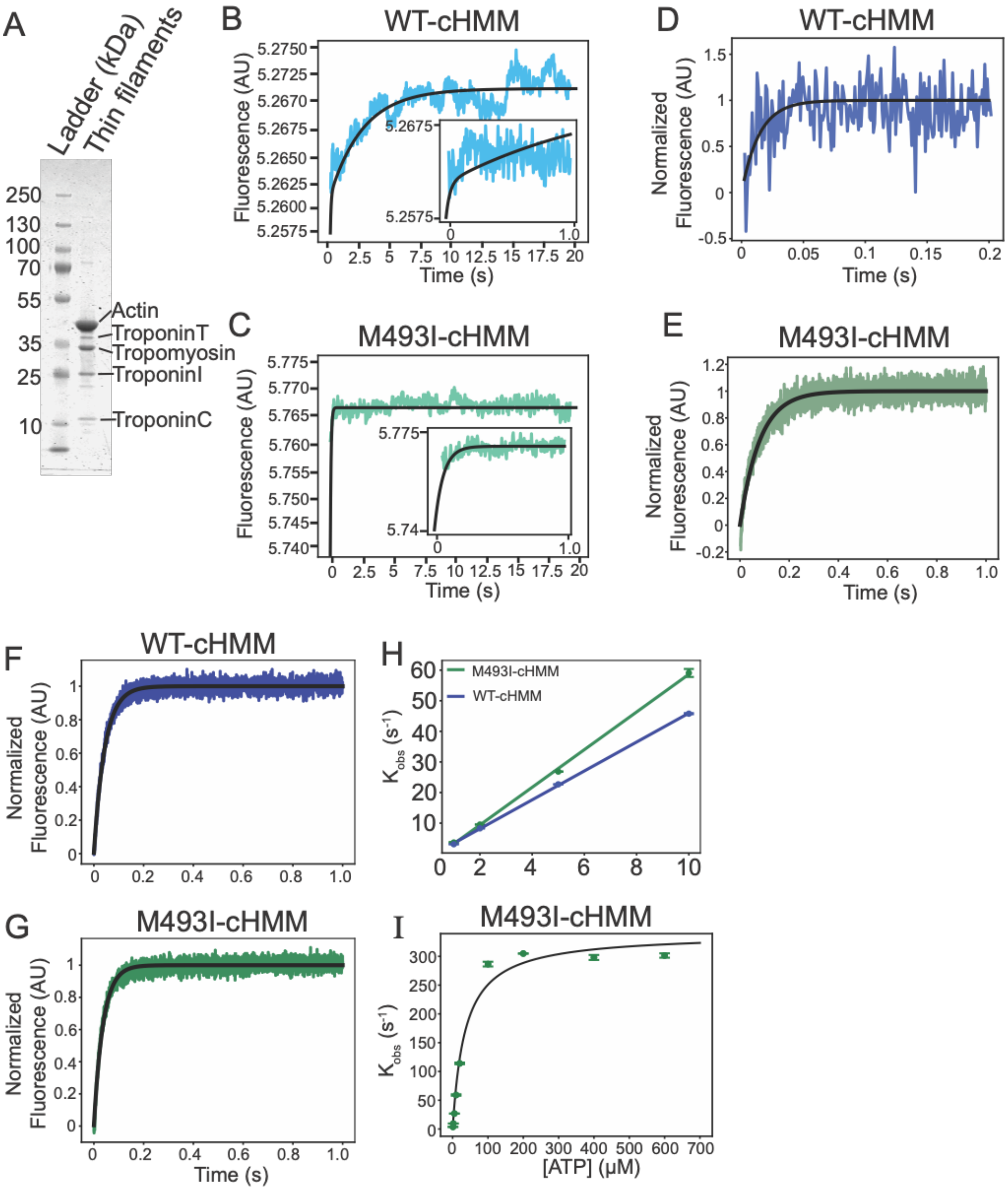


Fig S2: Transient kinetics of WT- and M493I-cHMMs. A) SDS-PAGE of porcine ventricular thin filaments (TFs). B-C) Example phosphate release transient of WT (B) with double exponential fitted curve and M493I (C, single exponential fit) sampled at 50 Hz. Insets: First 1 second of each trace, sampled at 250 Hz, with double exponential (WT) or single exponential (M493I) fits. D-E) Example transients of ADP release for WT (D) and M493I (E) from pyrene-labeled actin. F-

G) Example transients of ATP binding for WT (F) and M493I (G) at 5 µM ATP H) $K_{obs}$ for myosin dissociation by pyrene fluorescence change at low ATP concentrations to compare apparent 2$^{nd}$-order rate constant of ATP binding for M493I (green) *vs*, WT (blue) cHMM. Mean +/- S.D. as error bars. I) Michaelis-Menten curve fitted to dissociation rate *vs*. [ATP] for M493I.

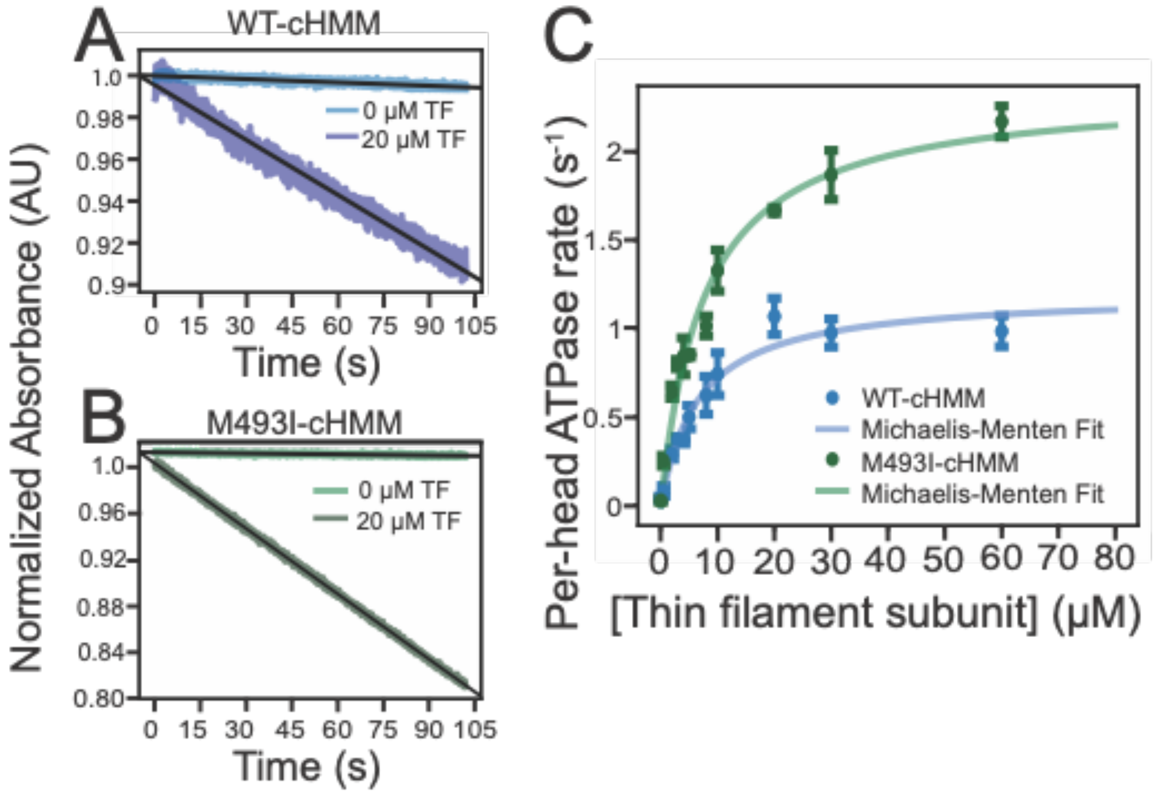


Fig S3: Actin-activated steady-state ATPase activity. A-B) Sample NADH absorbance traces from WT (A) and M493I (B) demonstrating linear decrease in the absence and presence of 20 μM TF subunit concentration. C) ATPase rate per head *vs*. [TF] with fitted Michaelis-Menten curves demonstrating approximate doubling of steady-state $V_{max}$ for M493I myosin relative to WT.

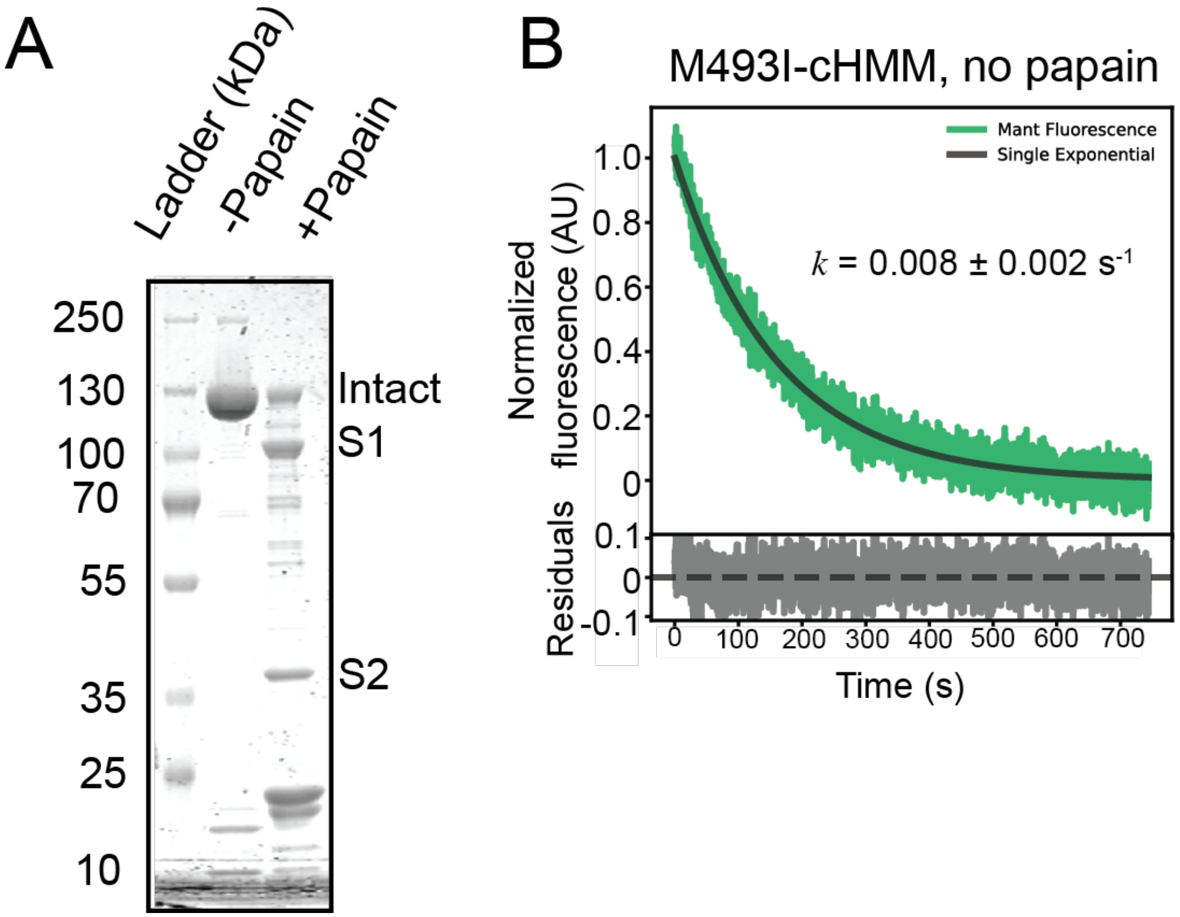


Fig S4: Papain digestion. A) SDS-PAGE of M493I myosin demonstrating cleavage into S1, S2, and limited intact myosin in the papain digest. B) Mant nucleotide fluorescence transient of an undigested M493I control using digestion buffer with E-64 papain inhibitor is not significantly different from that of M493I protein not exposed to the digestion conditions (Fig 4A).
